# Freshwater snail faecal metagenomes reveal environmental reservoirs of antimicrobial resistance genes across two continents

**DOI:** 10.1101/2025.02.26.640299

**Authors:** Angus M. O’Ferrall, Alexandra Juhász, Sam Jones, Peter Makaula, Gladys Namacha, Shaali Ame, David Oguttu, Aidan Foo, Sekeleghe A. Kayuni, James E. LaCourse, Janelisa Musaya, J. Russell Stothard, Adam P. Roberts

## Abstract

The transfer of antimicrobial resistance genes (ARGs) from environmental microbes to pathogens is a critical but underexplored One Health driver of antimicrobial resistance (AMR). Here, we evaluate freshwater snails, which are geographically widespread aquatic invertebrates, as environmental reservoirs of ARGs. We collected faeces from eight gastropod genera at 15 freshwater locations across Malawi, Uganda, Zanzibar, and the United Kingdom, and conducted the first freshwater snail faecal metagenomics study. We detected putative ARGs predicted to confer resistance to 13 antibiotic classes, including carbapenems in all countries. All ARGs that could be assembled into metagenome-assembled genomes (MAGs) were found within Proteobacteria, which dominated the faecal microbiomes and were strongly associated with the total ARG load. In Malawi, we linked *bla*_OXA-181_ (*bla*_OXA-48_-like), a previously mobilised carbapenemase gene, to *Shewanella xiamenensis*, the gene’s known progenitor. We detected another *bla*_OXA-48_-like gene by read-mapping from a sample in the United Kingdom. We identified mobile colistin resistance (*mcr*)-like genes at 11 of 15 locations, with two *mcr-7*-like genes being found in an *Aeromonas jandaei* MAG in Uganda. Our findings highlight freshwater snail faeces as a One Health-relevant environmental reservoir of clinically important ARGs.

**Data Summary:** Short reads from all samples sequenced in this study have been deposited in the Sequence Read Archive (SRA) under BioProject PRJNA1211045, with accession numbers SRX27371064 – SRX27371078. Data and code used to carry out analyses in R are available at https://github.com/amoreo71/freshwater_snail_faecal.

## Introduction

Antimicrobial resistance (AMR) is a growing global health emergency driven by industrial scale use of antimicrobials in medicine, veterinary practice, and agriculture since their discovery in the twentieth century (1). The impact of AMR on human health is largely driven by the acquisition of antimicrobial resistance genes (ARGs) in pathogenic bacteria (2). Consequently, most data used to investigate AMR originate from disease-causing organisms, yet there is increasing recognition that One Health frameworks accounting for the interdependence of human, animal, and environmental health systems need consideration to better understand the origins, drivers, and spread of AMR (3). This approach is essential for assessing how AMR may be linked to crossing planetary boundaries, such as those related to freshwater use, biodiversity loss, and climate change (4,5).

Outside the context of AMR, freshwater gastropod snails are already of interest in One Health research. Snails are globally distributed in aquatic environments, often playing critical roles as obligate intermediate hosts in the transmission of neglected tropical diseases when freshwater sources are contaminated with human and animal excreta. For instance, snails of the *Bulinus*, *Biomphalaria*, and *Oncomelania* genera are key in the transmission of schistosomiasis (6), while *Lymnaea* snails transmit fascioliasis (7). Recent efforts to characterise the microbiomes of medically important freshwater snails are generally motivated by the growing recognition that host-associated microorganisms can significantly influence host-parasite interactions (8). The microbiomes of *Biomphalaria* snails have been shown by 16S taxonomic profiling to be diverse and distinct from those of their freshwater environments (9), yet bacteriome transplant experiments have demonstrated the potential for these microbiomes to change depending on the surrounding environmental bacteria (10). Freshwater snail populations are highly mobile (11), capable of translocating due to changes in ecosystems caused by flooding or the formation of artificial water bodies (12,13), and are distributed across a range of human-influenced areas (14,15). Considering that the environments where freshwater snails graze— typically on detritus—can be contaminated with AMR enteric bacteria at sites of interaction with humans and animals, we wanted to explore the potential role of snails as widespread environmental reservoirs of ARGs (16). This may be particularly relevant in Low- and Middle-Income Countries (LMICs), where AMR is disproportionately linked to increased mortality (17), and where poor environmental health infrastructure is linked to AMR transmission across various One Health compartments (18,19).

The analysis of shotgun metagenomic sequence data is increasingly being used in the study of complex microbial samples and AMR, made possible by advances in next-generation sequencing and computational science technologies (20). Metagenomic studies are not reliant on culture steps that alter microbial communities and are therefore able to yield non-selective data on sample composition (21). Metagenomic analyses generally utilise short-read sequence data, which when coupled with the complexity of microbiome samples poses challenges in the assembly of metagenome-assembled genomes (MAGs) for low-abundance organisms without ultra-deep sequencing (22,23). Approaches that map reads directly to reference databases are therefore useful for taxonomic profiling and the detection of low-abundance target genes as they do not rely on metagenome assembly, and have proved to be more sensitive than assembly-based approaches for the detection of ARGs in complex samples such as faeces (24). When combining read-based profiling with assembly-based approaches that uncover the genomic context of AMR, allowing the linking of ARGs to specific hosts and mobile genetic elements (MGEs), metagenomics can enable a comprehensive understanding of the resistome and its potential impacts on microbial ecology and global health.

Here, we present a dataset characterising the first freshwater snail faecal metagenomes from 15 locations within Malawi, Uganda, Zanzibar (a self-governing region within the Republic of Tanzania), and the United Kingdom. Acknowledging that freshwater snail microbiomes vary by organ (25), and to mirror the natural mechanisms of faecal bacterial transfer from gastropods to the environment, we sampled faeces excreted by snails immediately after they were collected from their habitats, rather than extracting DNA directly from snail organs or rearing snails under laboratory conditions prior to DNA extraction. We use whole-genome metagenomics to investigate the role of freshwater gastropods as reservoirs, and potentially vectors, of AMR. Our goal is to identify clinically relevant ARGs within these common aquatic invertebrates and to examine their genomic context, providing insights and targets for future studies on ARG transfer to human and animal pathogens. We utilise direct read mapping for taxonomic profiling and ARG identification, followed by less sensitive but highly insightful assembly-based methods to explore the genomic context of ARGs.

## Methods

### Sampling, processing, and sequencing

In 2023 and 2024, we collected freshwater snails at 15 freshwater locations in four administrative regions across two continents, targeting snails of a single genus at each location (*Supplementary Table 1*). In Africa, collections were made from six locations in Malawi (five *Bulinus* and one *Cleopatra*), three locations in Uganda (one each of *Biomphalaria*, *Bulinus*, and *Physella*), and four locations on Unguja Island, Zanzibar (one each of *Bulinus*, *Cleopatra*, *Lanistes*, and *Pila*). In Europe, two locations in the United Kingdom were sampled (one *Anisus* and one *Lymnaea*). Sampling was opportunistic in nature and was carried out in conjunction with ongoing parasitological surveys that were not originally designed for balanced ecological comparisons between snail genera. As such, the presence of snail genera across the 15 sites reflects natural distributions at the time of sampling, rather than a standardised genus-level survey.

At each location, groups of 50 snails from the identified genus were gathered, rinsed, and temporarily pooled in sterile water. After 24 hours, faecal material deposited by each pooled snail group was collected to obtain adequate biomass for DNA extraction and sequencing. DNA was extracted from PBS-washed faecal samples with the QIAamp PowerFecal Pro DNA Kit (QIAGEN, Germany), as per manufacturer’s instructions. Concentration, purity and integrity of extracted DNA was assessed using a Qubit 4 fluorometer (Thermo Fisher Scientific, USA), NanoDrop One spectrophotometry (Thermo Fisher Scientific, USA), and the TapeStation System (Agilent Technologies, USA). Purified DNA extracts were shipped to Genewiz/AZENTA (UK) for library preparation, validation, and shotgun metagenomic sequencing. Sequencing was carried out with the Illumina NovaSeq instrument using a 2 x 150 bp paired-end configuration. The number of 150 bp Illumina paired end reads per sample ranged from 9,340,457 to 35,991,622 (*Supplementary Table 2*). The resulting 15 metagenomes proceeded to downstream analyses, each representing the faeces of 50 snails from a specific genus at a specific location.

### Bioinformatics

#### Read processing

Raw FASTQ sequence files were quality assessed using FASTQC (https://github.com/s-andrews/FastQC) Version 0.11.9. Trimming with Trimmomatic (26) Version 0.39 was performed to remove adaptor sequences and low-quality bases with a sliding window quality cutoff of Q20 and a minimum read length of 50 bp. Standard practice in metagenomic studies is to filter out host DNA by precisely aligning all reads to host reference genomes. As our study involved the analysis of pooled DNA samples from hosts whose genomes are not all available in databases, we applied the Kraken Software Suite protocol outlined by Lu et al. (27) to identify and extract contaminant and host DNA in a multi-step approach. Trimmed reads were classified using Kraken 2 (28) (Version 2.1.3). Kraken 2 works by matching read k-mers against a database of k-mers from existing genomes, before assigning each read to the lowest common ancestor (LCA) of all genomes that share those k-mers. We initially classified reads using the Kraken 2 standard database, which contains RefSeq archaea, bacteria, viruses, plasmid complete genomes, UniVec Core and the human reference genome, GRCh38. We then filtered out reads identified as eukaryotic (mapping to the human reference genome, either due to low-level human DNA contamination or because they were eukaryote-generic reads originating from snails) using the KrakenTools “extract_kraken_reads.py” script (https://github.com/jenniferlu717/KrakenTools). Retained reads were considered to have passed quality control.

We also built a custom Kraken 2 Gastropoda database containing the available GenBank snail genomes for all species within the genera represented in our samples: *Anisus vortex* (GCA_949126835.1), *Anisus vorticulus* (GCA_964264155.1), *Biomphalaria glabrata* (GCA_947242115.1), *Biomphalaria pfeifferi* (GCA_030265305.1), *Biomphalaria straminea* (GCA_021533235.1), *Biomphalaria sudanica* (GCA_036873155.1), *Bulinus truncatus* (GCA_021962125.1), *Lanistes nyassanus* (GCA_004794575.1), *Lymnaea stagnalis* (GCA_964033795.1), and *Physella acuta* (GCA_028476545.3). Genomes from the *Cleopatra* and *Pila* genera were not included, as they were not available within GenBank. Separately from classifications using the Kraken 2 standard database, we mapped all trimmed reads passing quality control to the custom-built Gastropoda database. We then extracted reads identified to be of gastropod origin, that had not already been discarded after being labelled as eukaryotic by the Kraken 2 standard database. This conservatively extracted subset of remaining Gastropoda reads that passed quality control (0.09 – 0.44% per sample) also underwent ARG detection steps to provide us with reassurance that our inability to precisely align each sample’s reads to a specific host reference genome did not bias ARG profiles.

#### Taxonomic profiling

Kraken 2 reports containing bacterial read taxonomic assignments were used to run species abundance estimation with Bracken (29) (Version 2.9), with a minimum number of reads required for a classification at the specified rank set at 10, to reduce noise from potential low-abundance contaminant species, as per default recommendations.

#### ARG database and clustering

We used the ResFinder database (30) (Version 2.3.2) to screen for ARGs. This database was selected because it focuses on acquired ARGs with known clinical relevance and mobile potential, thereby aligning with our aim to identify resistance mechanisms that may contribute to the spread of AMR from environmental reservoirs to clinical settings. Sequences in the ResFinder database were downloaded, then clustered using CD-HIT (31) (Version 4.8.1) to identify ARG clusters containing sequences with a minimum nucleotide identity of 90%, referred to as Cluster90s (24).

#### Read-based ARG detection

To identify ARGs within each sample, reads were aligned to reference sequences in the ResFinder database using KMA (32) (Version 1.4.14), a tool designed to map raw reads directly against redundant databases such as ARG databases. Alignments were then filtered to retain only those with a minimum 90% query identity and 60% template coverage—a common approach in the analysis of complex environmental metagenomes that balances accuracy and sensitivity by favouring reliable ARG identification while allowing for partial matches, thereby reducing the risk of missing low-abundance or divergent ARGs (33). For all reference sequences with alignment hits after filtering, the corresponding Cluster90 was identified and annotated manually. We calculated the normalised abundance of each ARG from read-mapping data, expressed as RPKM (Reads Per Kilobase per Million mapped bacterial reads), then summed the RPKM of all ARGs to calculate total ARG load per sample. We also performed read-based ARG detection on random sub-sets of reads (5 million, 10 million, 15 million, 20 million) from samples with > 20 million read pairs following quality control (Uk1, Z1, Z3, Z4). The ARG detection data generated from these read sub-sets was used to plot rarefaction curves and assess diversity saturation.

#### Metagenome assembly, binning, and classification

Reads identified as being bacterial by Kraken 2 were assembled with metaSPAdes (34) (Version 3.11.1), using the default k-mer spectrum (21, 33, 55). Assembly statistics were calculated with QUAST (35) (Version 5.0.2) using contigs ≥ 500 bp, to ignore low-quality sequences and ensure compatibility with ARG-containing regions (*Supplementary Table 2*). Contigs were indexed using the Burrows Wheeler Aligner (BWA) (Version 0.7.17), before bacterial reads were aligned to the indexed contigs with “bwa-mem” (36). The contigs and resulting sorted BAM files were parsed to the “jgi_summarize_bam_contig_depth” script from MetaBAT2 (37) (Version 2.17), before the resulting depth files were used by MetaBAT2 to bin assembled contigs of ≥ 2,500 bp to putative genomes (i.e., MAGs). CheckM (38) (Version 1.1.2) was used to assess the completeness and contamination of each MAG. CheckM outputs were compared to standards set by the genome standards consortium to classify MAGs as high-quality (>90% completeness, <5% contamination), medium-quality MAGs (>50% completeness, <10% contamination) or low-quality (<50% completeness and/or >10% contamination) (39). MAGs deemed as medium- or high-quality proceeded to further analysis. MAGs underwent taxonomic classification using GTDB-Tk (40) (Version 2.1.1) against the Genome Taxonomy Database (Version 09-RS220).

#### Assembly-based profiling of ARGs and MGEs

Assemblies were screened against the ResFinder (30) and ISfinder (41) databases using ABRicate (https://github.com/tseemann/abricate) (Version 1.0.1) at default settings (minimum 80% identity and 80% query cover) to identify contigs containing ARGs and IS elements. Compared to our read-mapping approach, in which we used a higher identity threshold to identify hits (90%) in combination with a lower minimum query cover (60%) due to the fragmented nature of using raw sequences to identify low-abundance genes, the assembly-based approach allowed for the detection of ARGs and MGEs that could then be characterised further within their genomic context following annotation. Annotations were performed using Bakta (63) (Version 1.9.3; full database Version 5.1). To elucidate the potential origins and dissemination pathways of ARGs, selected contigs were subjected to a sequence similarity search using the Nucleotide Basic Local Alignment Search Tool (BLAST) webtool (megablast algorithm) against the GenBank Nucleotide database. This step aimed to identify existing homologous sequences and/or gene clusters. The Clinker webtool (https://cagecat.bioinformatics.nl/tools/clinker) (64) was used to visualise gene cluster comparisons.

### Statistical analysis and visualisation

Species abundance and read-based ARG assay data were combined with hierarchical/classification data and sample metadata to generate a TreeSummarizedExperiment data container (65). Data associated with MAGs were handled separately. Within the R environment, downstream data wrangling, analysis, and visualisation took place predominantly using the R/Bioconductor *mia* and *miaViz* (Version 1.12.0) packages, and the R/CRAN *ggplot2* (Version 3.5.1) and *pheatmap* (Version 1.0.12) packages.

We calculated alpha diversity statistics at the species level, leveraging the high taxonomic resolution of shotgun metagenomics data. A Bray-Curtis dissimilarity matrix containing pairwise dissimilarities among microbiome samples was generated at species level using relative abundance data to investigate beta diversity and to perform principal coordinates analysis (PCoA). Ahead of hierarchical clustering of samples and taxa at the order level, centered log-ration (CLR) transformations were applied to microbial composition data (66) to normalise for compositional effects, with a pseudocount added to manage zero values using the *mia* package. Hierarchical clustering of both taxa and samples was applied to standardised CLR-transformed (CLR-z) data using Euclidean distance and complete linkage, as implemented by default in the *pheatmap* package. Before plotting ARG Cluster90 heatmaps, a log-transformation was applied to ARG load to enable visualisation across a wide range of abundances, with a pseudocount added to manage zero values using the *mia* package.

To assess monotonic relationships between numerical variables, such as the relative abundance of Proteobacteria and ARG load, we used Spearman’s rank correlation coefficient testing. This non-parametric test was chosen due to the small number of locations sampled (n = 15), its suitability for non-normally distributed data, and its robustness to outliers. Significance was determined with a threshold of *p* < 0.05. These analyses were followed by analyses of ARGs and MGEs found in MAGs, which supplement read-based profiles with high-resolution, species-specific context. Here, we used Wilcoxon rank-sum tests (non-parametric) to assess differences in the counts of ARGs and IS elements detected per complete MAG (calculated using raw ARG and IS element counts alongside genome completeness values per MAG) between taxa. Again, the threshold for statistical significance was set at *p* < 0.05.

## Results

### The freshwater snail faecal microbiome is diverse, varied, yet consistently rich in Proteobacteria

Species abundance estimations showed that Proteobacteria was the most abundant phylum observed, with a mean relative abundance of 66.27% (*Figure 1a*). Enterobacterales, an order within the Proteobacteria containing multiple species of clinical concern due to their ability to acquire and exchange ARGs, were well represented, with a mean relative abundance of 4.79%. The Uk1 sample was a notable outlier in which the relative abundance of Enterobacterales was estimated to be 41.70%. When excluding Uk1, the mean relative abundance of Enterobacterales dropped to 2.15%. Four orders were more abundant than Enterobacterales (Burkholderiales, Rhizobiales, Aeromonadales, and Pseudomonadales) all of which had a mean relative abundance of greater than 5% (*Figure 1b*). All samples exhibited high community (alpha) diversity. Shannon’s diversity index ranged from 3.881 (M5) – 7.372 (Z2) (mean = 6.474, median = 6.888). Simpson’s diversity index, in which scores from 0 – 1 are possible (1 = maximum possible diversity), ranged from 0.793 (M5) – 0.999 (Z2) (mean = 0.972, median = 0.994). We observed that alpha diversity (Shannon’s diversity index) was negatively corelated with the relative abundance of Proteobacteria (Spearman’s rank correlation coefficient = -0.636; *p* = 0.0129).

**Figure 1.**
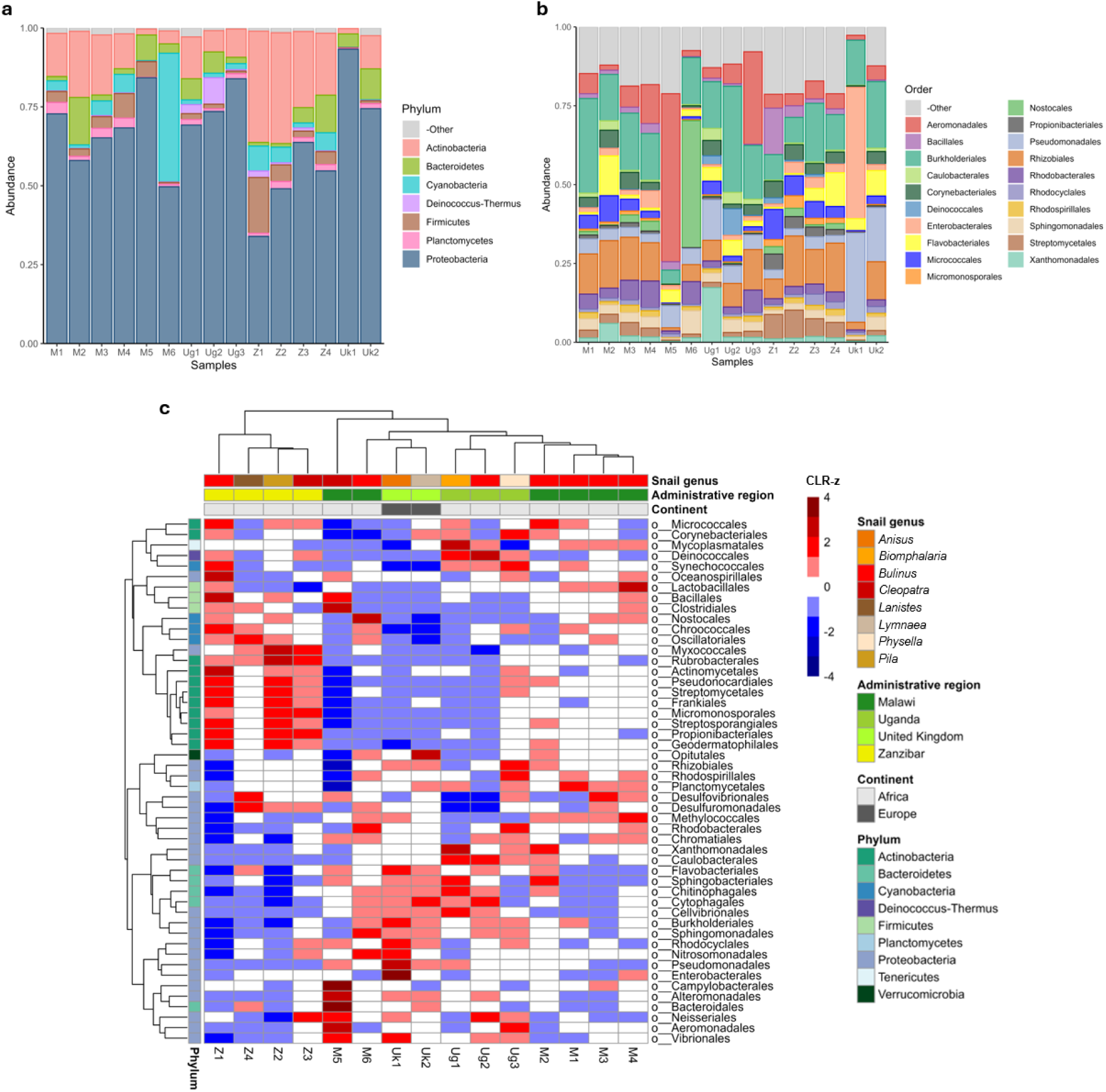
Exploration and comparison of bacterial taxa within freshwater snail faeces: a) Relative abundance of the top phyla; b) Relative abundance of the top orders; c) Clustered heatmap of standardised order abundances (CLR-z) across snail hosts and geographic locations. Column and row trees cluster samples and taxa, respectively, based on a hierarchical clustering approach. Taxa with a mean relative abundance < 1% are classified as ‘Other’ in parts a) and b).

Relative abundance data were aggregated at the order level, subjected to centred log-ratio (CLR) transformation, and subsequently standardised (CLR-z) to facilitate visual comparisons between taxa with varying absolute abundances (*Figure 1c*). From the 160 bacterial orders identified across all samples, the 50 most abundant orders, accounting for 97.92% of the total microbiome, were selected for hierarchical clustering and visualisation. Samples from the same administrative region clustered closely together, with the exception of Malawian samples, which were split into two distinct groups. This split corresponded to the two districts where the samples were collected: samples M1, M2, M3, and M4, collected in Mangochi District on the southern shorelines of Lake Malawi, clustered together, while samples M5 and M6, collected in Nsanje District, over 275 km from Mangochi District sampling sites, formed a separate cluster. Although they clustered together, northern hemisphere samples (United Kingdom) did not form a highly divergent cluster despite likely differences in climate-related and environmental conditions. Microbiomes from *Bulinus* snails (the most-represented genus, obtained from seven locations) showed notable divergence: the sample collected in Zanzibar clustered at one extreme of the dendrogram alongside samples from other snail genera, distinct from *Bulinus* samples collected in Uganda and the two Malawian districts.

When investigating diversity between samples at the species level, a Bray-Curtis dissimilarity matrix *(Supplementary Figure 1a)* showed a range of divergence between Malawian *Bulinus* snails. We observed moderate divergence (0.455) between two samples (M2, M4) collected a little over five kilometres from each other, both from stagnant water bodies close to the Lake Malawi shoreline, suggesting that local factors play a key role in influencing the microbiome. Despite these local variations, PCoA performed using Bray-Curtis distances at the species level *(Supplementary Figure 1b)* revealed similar locational trends to our order-level hierarchical clustering visualisation, in particular showing that the four samples from Mangochi District, Malawi (M1 – M4), as well as the four from Unguja Island, Zanzibar (Z1 – Z4), clustered closely together within their respective groups.

### ARGs conferring resistance to first- and last-line antibiotics detected by read mapping

Using read-based k-mer alignment (KMA) to screen for ARGs against the ResFinder database (30,32), reads mapped to 54 unique ARGs in total (minimum 90% query identity and 60% template coverage), belonging to 41 ARG Cluster90s (in which reference genes share minimum 90% nucleotide identity) (24). ARG richness (the number of unique ARGs identified by read mapping in any given sample) ranged from 1 – 16 (mean = 7.07, median = 6). ARG load ranged from 3.16 – 357.04 RPKM (mean = 87.99, median = 59.62). Although the depth of sequencing and number of unique ARGs detected per sample were not correlated (Spearman’s rank correlation coefficient = 0.283; *p* = 0.3065) (*Supplementary Figure 2*), we used random sub-sets of 5 million, 10 million, 15 million, and 20 million read pairs from samples with more than 20 million pairs passing quality control (Uk1, Z1, Z3, Z4) to further explore ARG saturation. Rarefaction curves showed that while ARG saturation can be reached by 10 million reads (e.g. Uk1), other samples (Z1, Z3, Z4) required up to 20 million reads to achieve saturation (*Supplementary Figure 3*). The ARGs that were not detected in smaller read sub-sets were only detected at very low in abundance in full read sets, therefore having a minimal effect on ARG load.

When we aggregated ARG relative abundance by class (i.e., the antibiotic that the ARGs confer resistance to), beta-lactamase genes (excluding carbapenemases) were the most abundant, contributing to 53.58% of the total ARG load across samples, followed by genes conferring resistance to colistin (14.38%), carbapenems (10.27%), sulfonamides (5.32%), tetracyclines (5.12%), aminoglycosides (2.77%), fosfomycin (2.15%), macrolides (2.14%), trimethoprim (1.90%), lincosamides (0.51%) chloramphenicol (0.47%), fusidic acid (0.19%), and quinolones (0.03%) (*Supplementary Figure 4*). Efflux pump genes conferring multidrug resistance contributed 1.16% of the total ARG load. Beta-lactamase genes from the OXA-12 Cluster90 were the most prevalent in our samples and were detected in 14 of the 15 metagenomes (*Figure 2*). We also detected ARGs from the CphA Cluster90, a family of carbapenem-hydrolyzing metallo-beta-lactamase genes, in 10 of the 15 samples. Three OXA-carbapenemase genes were identified, each from a different country. These included two ARGs from the OXA-48 Cluster90 (*bla*_OXA-181_ and *bla*_OXA-547_, in Malawi and the United Kingdom, respectively), of concern globally due to plasmid-borne dissemination amongst Enterobacterales (46), and *bla*_OXA-212_ (Uganda) from the OXA-211 Cluster90 of carbapenemase genes that naturally occur in *Acinetobacter johnsonii* (47). We also detected putative colistin resistance genes, predominantly variants of the *mcr-7.1* gene, in eleven metagenomes.

**Figure 2.**
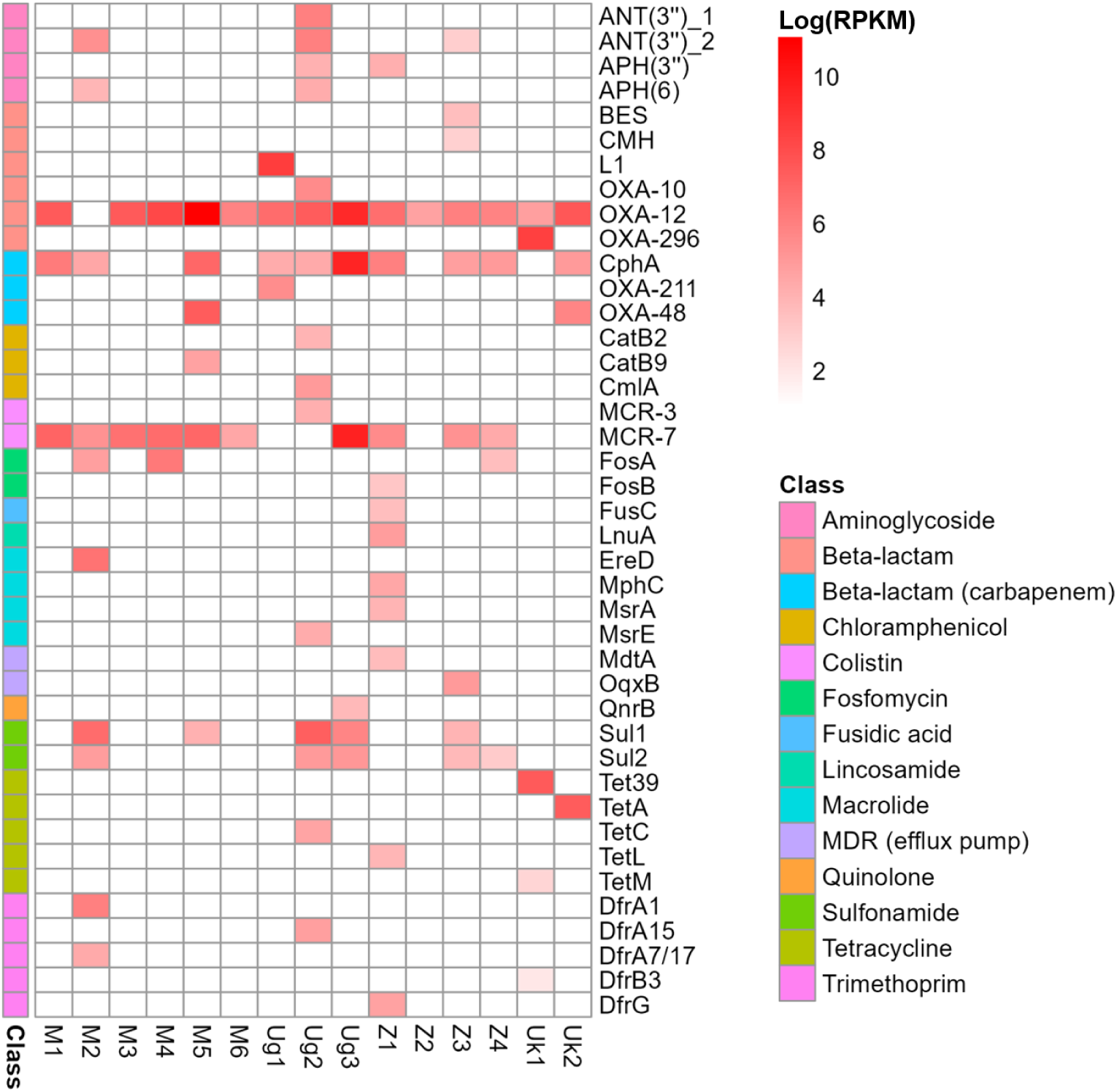
Heatmap of normalised ARG Cluster90 abundances across samples, expressed as log(RPKM). Cluster90s are colour-coded by the antibiotic class that they are predicted to confer resistance against.

We observed a positive correlation between the relative abundance of Proteobacteria (the predominant bacterial order) and ARG load (Spearman’s rank correlation coefficient = 0.846; *p* < 0.0001) (*Figure 3a*). A significant association between Proteobacteria relative abundance and ARG richness was not detected (Spearman’s rank correlation coefficient = 0.294; *p* = 0.2877). Among the 20 orders of bacteria with a mean relative abundance above 1% (*Supplementary Table 3*), only Pseudomonadales had a significant positive correlation with ARG load (Spearman’s rank correlation coefficient = 0.604; *p* = 0.0195), although this was not a clear linear trend (*Figure 3b*). Meanwhile, the relative abundance of Aeromonadales was particularly high in two samples (M5, Ug3), within which ARG load was also markedly higher (> 300 RPKM) than all other samples (< 100 RPKM). However, the overall correlation between samples was deemed non-significant (Spearman’s rank correlation coefficient = 0.357; *p* = 0.1916). These results indicate that different orders within the Proteobacteria are involved in AMR to varying extents in different snail microbiomes, and that assembly-based methods that elucidate the genomic context of AMR are needed to explore this with greater resolution.

**Figure 3.**
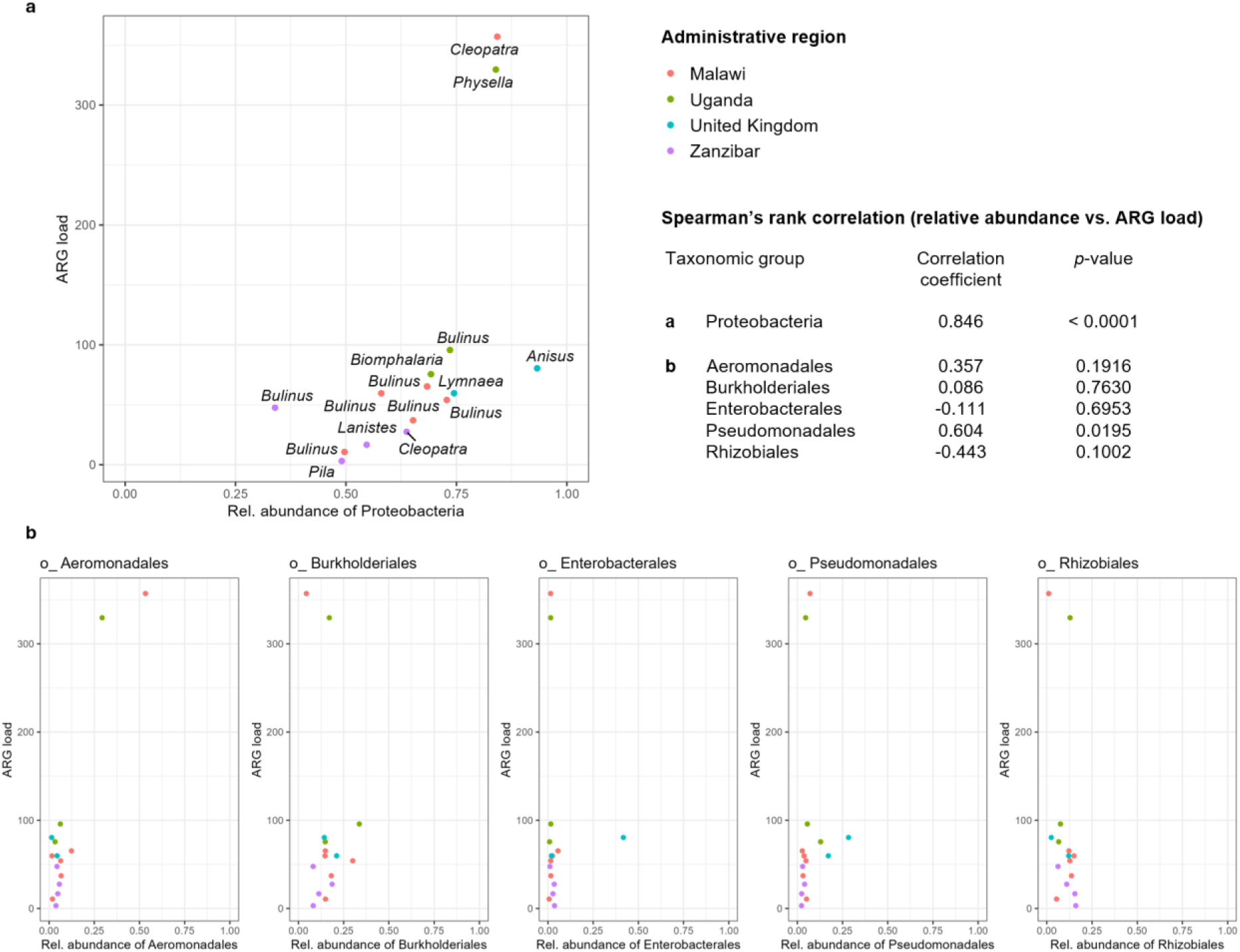
Associations between Proteobacteria and ARG load: a) Relative abundance of Proteobacteria versus ARG load; b) Relative abundance of the five most abundant orders within the Proteobacteria versus ARG load. Within-plot Labels in part a) are snail genera. Spearman’s rank correlation coefficients and associated p-values are presented for each taxonomic group for relative abundance versus ARG load.

## Assembly-based methods link ARGs and MGEs to various bacterial genera and identify known progenitors of mobile carbapenemase and colistin resistance

We detected fewer ARGs by screening metagenome assemblies against the ResFinder database using ABRicate (mean = 2.53, median = 2) than we did by read mapping (*Supplementary Figure 5*). However, we were able to acquire information on the carriage of ARGs by specific taxonomic groups following assembly and binning of contigs. Across all samples, 10 of the 38 ARGs (26.3%) assembled on contiguous sequences (contigs) could be binned to high-quality (>90% completeness, <5% contamination) or medium-quality (>50% completeness, <10% contamination) MAGs. These ARGs were all binned to MAGs classified as Proteobacteria (genera: *Aeromonas*, n = 7; *Shewanella*, n = 1; *Stenotrophomonas*, n = 2) (*Figure 4*). Specifically, these MAGs all belonged to the Gammaproteobacteria class, from which ARGs were identified at a rate of 0.62 per complete genome. Even within our small dataset of 29 medium- and high-quality MAGs, the difference in ARG detection rates in Gammaproteobacteria, which made up 85.7% (18/21) of Proteobacteria MAGs and 62.1% (18/29) of all MAGs, was significantly higher than in all other bacterial classes (*p* = 0.0391). We detected between 8 and 97 putative IS elements per sample (mean = 37.8, median = 29). IS elements were more commonly identified in Gammaproteobacteria MAGs (1.03 per complete genome) than in other MAGs (0.57 per complete genome). Although the difference observed did not meet the threshold for statistical significance within this relatively small dataset (*p* = 0.1223), the trend likely reflects underlying biological differences in MGE content between taxonomic groups.

**Figure 4.**
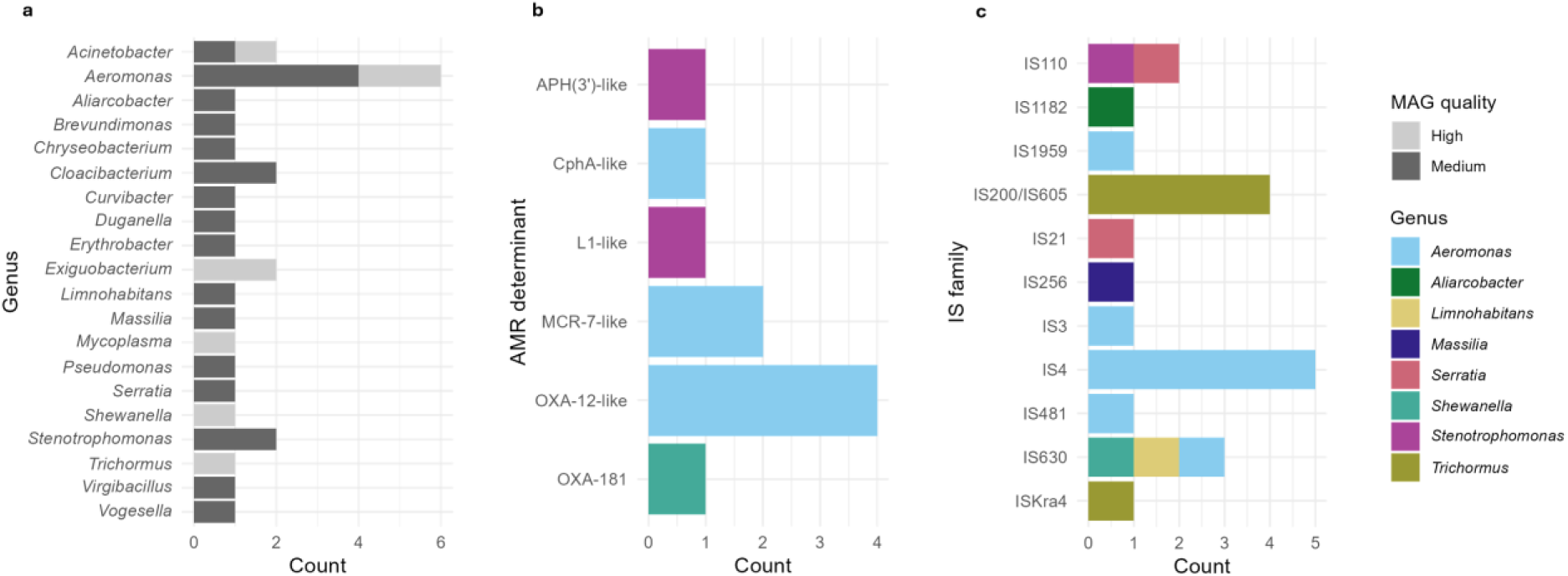
Summary of medium- and high-quality MAGs – a) Genera; b) AMR determinants (annotated as ‘-like’ if nucleotide identity is <100% when compared to reference); c) IS families.

We identified two contigs containing both IS elements and ARGs, neither of which could be binned to a MAG. A 4042 bp contig (Ug2 NODE_1302: *Bulinus* spp. faeces, Lake Victoria, Uganda) contained *sul1* (sulfonamide resistant dihydropteroate synthase) and *bla*_OXA-10_ (beta-lactamase) genes, on which both ARGs were located upstream of IS*6100*. The gene cluster also contained *qacEdelta* (a gene conferring resistance to antiseptics) and *yhbS* (putative N-acetyltransferase). The contig displays 100% query cover with one complete genome in the GenBank Nucleotide database—*Aeromonas hydrophila* (accession no. AP025277)—with 99.97% identity, but also maintains high query cover (> 95%) and identity (> 99%) with chromosomal and plasmid sequences from various other bacteria (*Pseudomonas aeruginosa*, *Vibrio cholerae*, *Morganella morganii*, *Salmonella enterica*, *Klebsiella pneumoniae*). A second, shorter, 3505 bp contig (M2 NODE_687: *Bulinus* spp. faeces, Mangochi District, Malawi), aligned with 99.94% nucleotide identity to Ug NODE_1302 over the region containing *qacEdelta, sul1* and *yhbS* upstream of IS*6100* (*Figure 5*), with a single nucleotide polymorphism (SNP) in the *sul1* gene. The similarity observed between the two contigs from Ug2 and M2, and with sequences from a range of bacterial species in GenBank, demonstrates the widespread nature of certain ARGs associated with MGEs, which we show can be found in freshwater snail faeces. However, we could only link MGEs directly to ARGs on contigs in the above two examples. Searches for ARGs associated with IS elements in transposons should ideally capture 24.34 kbp of DNA (based on the length of composite and unit transposons in The Transposon Registry) (48). The median length of contigs containing ARGs in our dataset was 2.10 kbp, demonstrating the limitations in assembling large contigs from short reads in diverse metagenomic samples (23), and the associated inability to capture all co-localised ARGs and MGEs.

**Figure 5:**
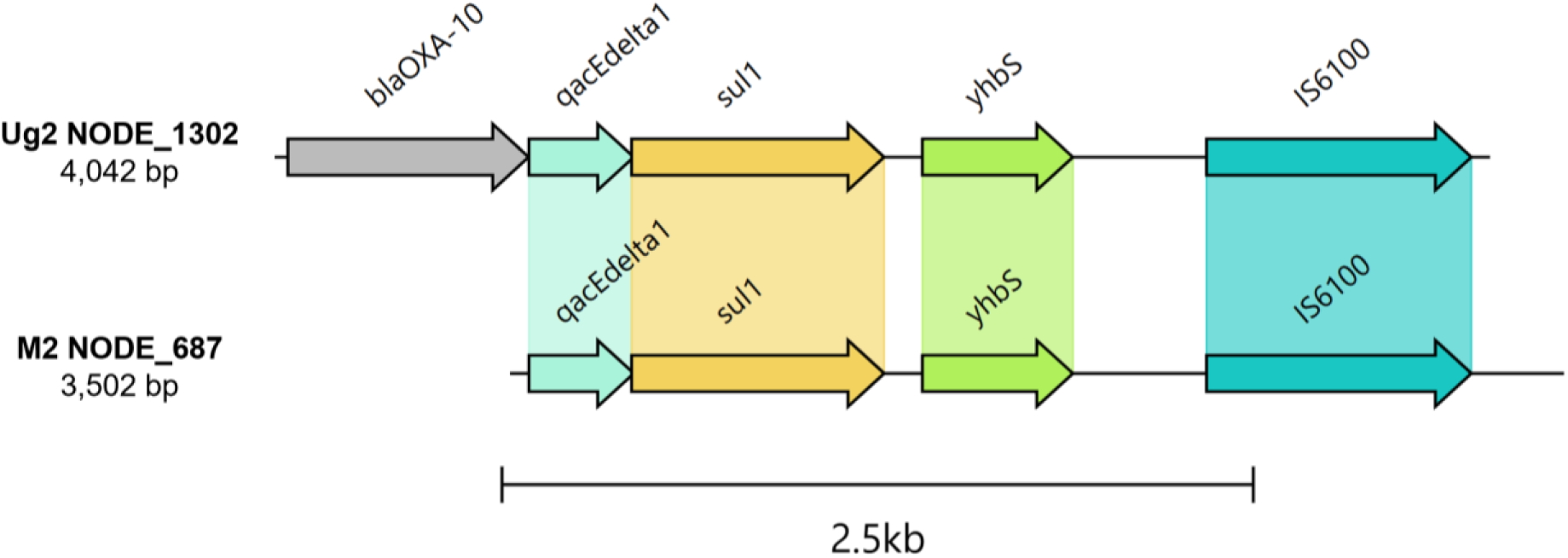
Gene cluster comparison between Ug2 NODE_1302 and M2 NODE_687 containing bla_OXA-10_ (Ug2 only), qacEdelta1, sul1 and yhbS upstream of IS6100. Regions coding for hypothetical proteins are not displayed.

One of the two *bla*_OXA-48_-like carbapenemase genes identified by read-mapping was also detected in metagenome assemblies. NODE_173 from the M5 metagenome (*Cleopatra* spp. faeces, Nsanje District, Malawi) contained *bla*_OXA-181_ on a 20,996 bp contig with 17 open-reading frames (ORFs). The contig was binned to a high-quality MAG classified as *Shewanella xiamenensis* (*Table 1*). When the *bla*_OXA-181_-containing contig was queried against the GenBank Nucleotide database, the closest alignment was to chromosomal DNA from an *S. xiamenensis* isolate (accession no. AP026732) obtained from urban drain water from Vietnam (100% query cover, 98.51% identity) (49). With only one SNP in a non-coding region, the contig also aligns with 99.92% identity over a 1309 bp region (8073–9381) to various plasmids from the Enterobacterales containing the internationally disseminated *bla*_OXA-181_ gene and a truncated *lysR* (Δ*lysR*) gene, including the KP3-A plasmid from which *bla*_OXA-181_ was first characterised in a clinical *K. pneumoniae* isolate (accession no. JN205800) (50). The region containing *bla*_OXA-181_ and Δ*lysR* within the KP3-A (and other) plasmid(s) has previously be traced back to its origin, *S. xiamenensis*, where *bla*_OXA-181_ is found adjacent to a complete *lysR* gene (51), as is also seen in our annotated sequences (*Supplementary Figure 6*). The ability to detect, assemble, and bin this critical priority ARG is aided by the high relative abundance of *Shewanella* in the M5 sample (5.07%). We also observed a strong *Shewanella* signal in the Uk2 sample (relative abundance = 1.33%), from which another *bla*_OXA-48_-like variant (*bla*_OXA-547_) was detected by read-mapping, when compared to the remaining 13 samples in which no *bla*_OXA-48_-like genes were detectable (mean *Shewanella* relative abundance = 0.11%).

**Table 1:**
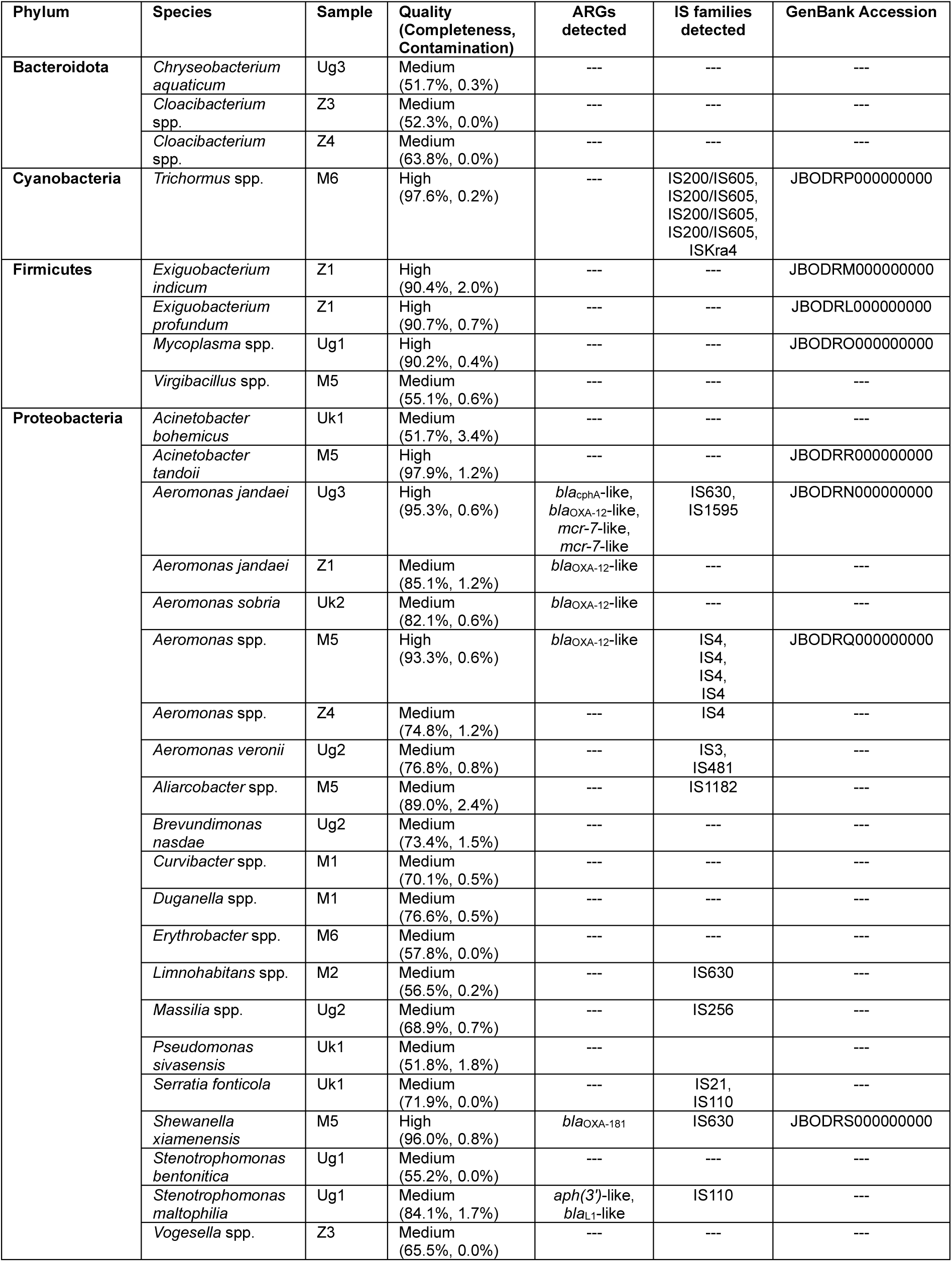
Classification of MAGs recovered from freshwater snail faecal microbiomes, with quality metrics, ARGs and IS elements detected, and GenBank Accession numbers for high-quality MAGs.

Two *mcr-7*-like lipid A phosphoethanolamine transferase genes were detected in a high-quality *Aeromonas jandaei* MAG (*Table 1*) recovered from the Ug3 metagenome (*Physella* spp. faeces, Kimi Island, Uganda) on separate contigs (NODE_28 and NODE_32: 86.88% and 81.93% nucleotide identity to *mcr-7.1*, respectively). Lipid A phosphoethanolamine transferase enzymes cause a reduction in overall net-negative charge of the outer membrane, conferring resistance to colistin among some gram-negative bacteria, and are a major public health concern due to their propensity to be mobilised from progenitor species chromosomes to plasmids, limiting last-resort antibiotic treatment options for infections caused by multi-drug-resistant pathogens (52). *Aeromonas* spp. are the known progenitors of *mcr-3* and *mcr-7* (53). From our 15 samples, reads mapped to the MCR-3 Cluster90 in one and the MCR-7 Cluster90 in 10, underscoring the widespread prevalence of *Aeromonas*-associated *mcr*-like genes in the sampled environment.

## Discussion

Our findings provide novel insights into the role that freshwater snails play as AMR reservoirs, revealing a potential widespread environmental source of clinically relevant ARGs. Through analysis of snail faecal metagenomes from diverse geographic locations, we demonstrate that their microbiomes harbour a wide array of bacterial species and ARGs, including those conferring resistance to both first- and last-line antibiotics. Identification of environmental progenitors of potentially mobile ARGs conferring resistance to carbapenems and colistin is particularly noteworthy in sub-Saharan Africa, where such antibiotics are not commonly available or prescribed for patients with severe infections caused by Enterobacterales, but where resistance among Enterobacterales to other treatment options such as penicillins, third-generation cephalosporins, and gentamicin has increased (54). A likely increase in carbapenem prescribing as a result of this may introduce additional selective pressures to aquatic habitats, driving selection and transfer of ARGs from environmental reservoirs to clinically relevant bacteria (55). Our results emphasise the potential for snails to contribute to this dissemination in freshwater ecosystems, particularly in those modified by anthropogenic change with deficits in local WASH (water, sanitation, and hygiene) infrastructures (3,19), although we are not able to confirm ARG mobility from these sequencing data alone.

Proteobacteria dominated the snail faecal microbiomes, reinforcing results from previous studies that found Proteobacteria to be highly abundant in other freshwater snail microbiomes such as the gut and haemolymph (9,56–58). The significant correlations observed between Proteobacteria abundance and both alpha diversity and ARG load suggest that an abundance of Proteobacteria reduces overall microbiome diversity while increasing the relative abundance of ARGs within the microbial community. We binned genes from the three most prevalent ARG Cluster90s (CphA, MCR-7, OXA-12) to *Aeromonas* spp. MAGs, highlighting this group of bacteria as geographically widespread AMR reservoirs. The Proteobacteria order and *Aeromonas* genus have both recently been shown to be core taxa in gut samples extracted from laboratory-reared *Biomphalaria* snails (58); as well as signalling that snail faecal microbiomes are a reflection of the intestinal tract, our results indicate that these core features are preserved across various snail genera and within their natural habitats. Meanwhile, our hierarchical clustering and PCoA visualisations demonstrate the variation in numerous other bacterial taxa between snails of the same genus (*Bulinus*) observed between countries, and between districts in the case of Malawi. The moderate diversity observed between proximal samples from Malawian *Bulinus* snails at the species level highlight the potential role of local environmental conditions, such as water pollution, climate, and antibiotic exposure, in addition to host-specific characteristics, in shaping faecal microbiomes beyond the core taxa.

We recovered eight high-quality MAGs and 21 medium-quality MAGs from 20 bacterial genera in total, yet 1246 unique genera were detected in total by Kraken 2 read profiling, demonstrating the limitations of assembly-based approaches in complex samples such as these. It is therefore expected that fewer ARGs were detectable in MAGs, which are generally recovered from high-abundance organisms, than by read mapping or by screening all contigs, including those that could not be binned to MAGs. The inability to recover medium- to high-quality MAGs (with >50% completeness) in 1226 of the 1246 genera detected by read profiling (98.39%) suggests that some ARGs from low-abundance species are likely to have been undetectable even by our read-based approach, particularly in samples with lower sequencing depths. We specified that 60% template coverage was needed to call a hit to reduce the chance of false positives, and hence the ARG profiles that we present here are likely biased towards those found in high-abundance genera such as *Aeromonas*. This idea is supported by our rarefaction analysis, in which varying sequencing depths were required to achieve ARG saturation. Nonetheless, ARGs associated with abundant bacterial taxa were commonly detected across the four samples with fewer than 10 million reads.

The detection of ARGs from the CphA, MCR-7 and OXA-12 Cluster90s in 66.7%, 66.7% and 93.3% of samples, respectively, all of which originate from *Aeromonas* species (59,60), is a reflection of the skew towards the detection of ARGs associated with abundant taxa. The *mcr-7.1* colistin resistance gene was first identified on plasmids in *K. pneumoniae* isolates from chickens in China (61), but chromosome-borne variants of the gene have more recently been identified in *Aeromonas* isolated from poultry in the United States (62), and in global environmental sources (water) (63). The suggestion that *Aeromonas* species are the natural reservoirs of *mcr-3* and *mcr-*7 was since proven by robust evolutionary models (53), and we were able to assemble and bin *mcr-*7-like variants to a putative *Aeromonas jandaei* MAG. Our results signpost to the potential for emergence of new variants of these ARGs from freshwater gastropod faeces, and specifically, the *Aeromonas* reservoirs within.

Similar reservoirs of ARGs in low-abundance species may be more widespread than our data indicate. We demonstrate the presence of *bla*_OXA-181_ and *bla*_OXA-547_ in samples from Malawi and the United Kingdom, respectively, both of which are *bla*_OXA-48_*-*like genes. In the case of *bla*_OXA-181_, we could bin the > 20 kbp contig containing the gene to a high-quality MAG of *S. xiamenensis*, the species known to be the gene’s progenitor (51). *Shewanella* species have been identified as the progenitors of numerous *bla*_OXA-48_ variants (64), and the enzymes coded for by these genes hydrolyse carbapenems and confer resistance to most beta-lactam, beta-lactamase inhibitor combinations. They emerged and proliferated over the last two decades to become the most prevalent carbapenemases amongst Enterobacterales in many parts of Europe, Northern Africa, and the Middle East (46). Consequently, they are a major public health concern, with carbapenemase-producing Enterobacterales (CPE) being listed by the World Health Organisation as critical priority pathogens for which new antimicrobial agents are urgently needed (65). While we cannot infer mobilisation of *bla*_OXA-48_- or *mcr*-like genes from environmental species to Enterobacterales in our study, the rapid dissemination of ARGs co-localised with IS elements among various species within the Enterobacterales often occurs as a result of one initial mobilisation event, after which transposable units can move between replicons and cells (64,66).

To our knowledge, this is the first report of *bla*_OXA-181_ in Malawi, a country where CPE are uncommon in the clinical setting, but extended-spectrum beta-lactamase (ESBL)-producing Enterobacterales resistant to a range of other beta-lactam antibiotics are, and contribute to increased mortality in hospital patients (67,68). ESBL-producing Enterobacterales have also been widely detected in water systems as well as human and animal faeces in Malawi (68,69). Efforts to characterise the selective pressures behind the spread of ESBL-producing Enterobacterales in Malawian wastewater and freshwater are ongoing, but low environmental antimicrobial concentrations are known to drive the transfer and persistence of ARGs in water systems (55). This selective pressure may also facilitate the emergence of *bla*_OXA-48_*-*like genes from environmental reservoirs where environmental and human health intersect. Meanwhile, such reservoirs of carbapenem resistance genes could allow bacteria to react rapidly if carbapenems are introduced more routinely into clinical practice, limiting their efficacy. The emergence and spread of ARGs is often driven by horizontal gene transfer (HGT) associated with MGEs, such as conjugative plasmids, which regularly carry multiple IS elements (3,70). We show that IS elements are abundant within snail faecal microbiomes. However, the suggestion of HGT is purely hypothetical in our context, without capturing widespread co-localisation of ARGs on MGEs and performing functional assays. We could only demonstrate co-localisation of ARGs and IS elements in two instances where *sul1*-containing gene clusters were identified upstream of IS*6100* in samples from Malawi and Uganda. This is a common issue in short-read metagenomic due to the fragmented nature of assemblies (71).

Our study does have several limitations. For example, the use of pooled snail faeces for metagenomic analysis, while necessary to obtain sufficient biomass for DNA extraction and sequencing, limits the ability to assess individual snail variability and the potential heterogeneity in microbiome composition and ARG content in snails from the same location. Meanwhile, the use of short-read sequencing, although effective for taxonomic profiling and ARG identification, poses challenges in assembling complete genes and genomes, particularly for low-abundance species, potentially leading to an underestimation of the full richness of ARGs. This limitation is compounded by the inherent bias toward detecting ARGs in more abundant taxa which may skew the representation of ARGs in less abundant microbes. Ultra-seep sequencing may be required to detect such ARGs, even by read mapping methods that do not rely on metagenome assembly. Additionally, while read-based ARG detection approaches are widely used in the analysis of complex metagenomes to enable the detection of low-abundance ARGs in the absence of assembly (24,33), partial hits may represent truncated or non-functional genes. Inherently, short-read metagenome assemblies from diverse samples are highly fragmented. This limited our ability to explore co-localisation of ARGs and on plasmids and composite transposons. The accuracy of, and access to, long-read metagenomics sequencing technologies is rapidly improving (71), and these technologies could be used in future metagenomic studies to address the issues associated with fragmented assemblies. Proximity ligation methods such as Hi-C could also assist by linking MGEs to host genomes (72), though this was not a feasible addition to our study due to limited sample biomass and the need for dedicated extraction protocols. Furthermore, while this study highlights the potential role of freshwater snails as AMR reservoirs, the cross-sectional nature of the sampling limits our ability to infer causality or the dynamics of ARG transmission over time. Longitudinal studies are needed to better understand temporal changes in ARG prevalence and the environmental factors influencing these patterns. Finally, while the study identifies ARGs predicted or known to be of clinical relevance, it does not explore the functional impact of these ARGs, nor does it confirm their potential for HGT to human pathogens, a critical future step to quantify the risks associated with our findings.

In conclusion, our study represents the first metagenomic analysis of AMR in freshwater gastropod faeces, providing important insights into the potential roles of these invertebrates as reservoirs of clinically relevant ARGs. Our findings have significant implications for AMR surveillance within One Health frameworks, particularly in areas where agricultural animals, wild animals, and humans utilise the same open water sources. This reservoir of ARGs poses a threat to public health in an era of industrial-scale antibiotic use, particularly where access to clean water and sanitation is limited. Future research should build on these results through longitudinal sampling and by investigating how environmental factors, such as the presence and concentration of antibiotics, influence ARG richness and abundance in snail microbiomes. Meanwhile, incorporating long-read sequencing, proximity ligation methods, and functional assays in future work, where feasible, will improve our understanding of ARG mobility. Additional sampling in nearby human and animal populations to map potential transmission of ARGs from snail-associated bacteria to human pathogens will be essential for designing targeted interventions to curb AMR spread from and into shared aquatic environments.

## AUTHOR STATEMENTS

### Author contributions

The study was conceived and designed by AMO’F, JM, JRS and APR. Sampling was carried out by AMO’F, AJ, SJ, PM, GM, SA, DO, SAK, JEL and JRS. Formal analysis was performed by AMO’F, AF, JRS and APR. The original draft was prepared by AMO’F, JRS and APR, then reviewed and edited by all authors. All authors approved the final version of the manuscript.

## Acknowledgements

The authors thank Dr Simon Wagstaff and Mr Andrew Bennet for provision and maintenance of the scientific computing environment at LSTM used to carry out bioinformatic analysis. The authors acknowledge staff and local people in each of the sampling areas who aided the sampling teams.

