## Supplementary materials for "Freshwater snail faecal metagenomes reveal environmental reservoirs of antimicrobial resistance genes across two continents"

*Supplementary Table 1: Sample characteristics.*

*Supplementary Table 2: Summary of Illumina sequencing data and metagenome assemblies.*

*Supplementary Table 3: Correlation between relative abundance of the top 20 bacterial orders and ARG load.*

*Supplementary Figure 1. Species-level beta-diversity: a) Bray–Curtis dissimilarity matrix; b) Principal coordinates analysis (PCoA) performed using Bray-Curtis distances.*

*Supplementary Figure 2: Scatter plots showing associations of sequencing depth with: a) ARG detection; b) MAG recovery.*

*Supplementary Figure 3: Rarefaction curves showing the number of unique ARGs detected following random sub-setting of reads from samples with > 20 million reads following trimming and filtering.*

*Supplementary Figure 4: Relative abundance of ARGs aggregated by class (RPKM).*

*Supplementary Figure 5: Read- versus assembly-based ARG detection.*

*Supplementary Figure 6: Annotation of M5 NODE_173 containing $bla_{OXA-181}$ and ΔlysR, compared to Tn2013 from K. pneumoniae strain KP3 plamid pKP3-A (Accession no. JN205800) containing $bla_{OXA-181}$ flanked by ΔlysR-ΔereA and ISEcp1.*

*Supplementary Table 1: Sample information and characteristics.*

| ID | Administrative region | Continent | Site name | Water type | GPS East | GPS South | Snail genus |
|---|---|---|---|---|---|---|---|
| **M1** | Malawi | Africa | Nkopola | Stream | -14.314 | 35.144 | *Bulinus* |
| **M2** | Malawi | Africa | Palm Beach | Stagnant | -14.392 | 35.221 | *Bulinus* |
| **M3** | Malawi | Africa | Mangochi Town | River | -14.452 | 35.242 | *Bulinus* |
| **M4** | Malawi | Africa | Chipereka | Stagnant | -14.385 | 35.275 | *Bulinus* |
| **M5** | Malawi | Africa | Mthawira | Stagnant | -16.854 | 35.302 | *Cleopatra* |
| **M6** | Malawi | Africa | Nsanje Port | Stagnant | -16.932 | 35.264 | *Bulinus* |
| **Ug1** | Uganda | Africa | Ngamba Island | Lake | -0.106 | 32.653 | *Biomphalaria* |
| **Ug2** | Uganda | Africa | Kimi Island 1 | Lake | -0.085 | 32.645 | *Bulinus* |
| **Ug3** | Uganda | Africa | Kimi Island 2 | Stagnant | -0.088 | 32.652 | *Physella* |
| **Z1** | Zanzibar | Africa | Bumbwini | Stream | -5.954 | 39.195 | *Bulinus* |
| **Z2** | Zanzibar | Africa | Donge Mbije | Stagnant | -5.930 | 39.256 | *Pila* |
| **Z3** | Zanzibar | Africa | Mwera 1 | Stagnant | -6.142 | 39.269 | *Cleopatra* |
| **Z4** | Zanzibar | Africa | Mwera 2 | Stagnant | -6.137 | 39.266 | *Lanistes* |
| **Uk1** | United Kingdom | Europe | Knowsley | Stream | 53.437 | -2.819 | *Anisus* |
| **Uk2** | United Kingdom | Europe | Preston Montford | Stream | 52.724 | -2.840 | *Lymnea* |

*Supplementary Table 2: Summary of Illumina sequencing data and metagenome assemblies.*

| ID | Illumina short reads | | | Bacterial assembly (contigs ≥ 500 bp) | | | Proportion of bacterial reads mapping to assembly (%) |
|---|---|---|---|---|---|---|---|
| | **Total read pairs** | **Read pairs passing QC** | **Bacterial read pairs after QC** | **Assembly size (bp)** | **Total contigs** | **N50** | |
| **M1** | 9,340,457 | 7,452,698 | 2,142,223 | 21,422,957 | 18,427 | 1,369 | 45.79 |
| **M2** | 12,257,171 | 10,083,776 | 3,342,252 | 49,476,907 | 52,617 | 952 | 57.66 |
| **M3** | 9,982,521 | 7,954,558 | 1,905,474 | 9,999,998 | 12,847 | 746 | 25.24 |
| **M4** | 9,629,854 | 7,647,992 | 890,800 | 5,267,659 | 7,129 | 712 | 15.70 |
| **M5** | 9,476,943 | 7,576,553 | 3,974,033 | 34,276,162 | 20,545 | 2,699 | 85.05 |
| **M6** | 12,399,711 | 9,917,053 | 3,993,286 | 38,011,353 | 36,921 | 1,000 | 70.24 |
| **Ug1** | 17,829,973 | 13,669,537 | 4,446,415 | 71,092,332 | 68,500 | 1,030 | 69.58 |
| **Ug2** | 18,211,831 | 14,031,764 | 5,008,151 | 67,825,877 | 65,357 | 1,081 | 75.54 |
| **Ug3** | 19,554,138 | 14,801,846 | 5,708,500 | 62,394,277 | 65,605 | 929 | 79.15 |
| **Z1** | 28,260,918 | 20,934,433 | 4,494,974 | 44,526,232 | 43,914 | 1,015 | 57.69 |
| **Z2** | 26,317,771 | 19,886,506 | 5,172,786 | 31,591,691 | 41,009 | 738 | 39.82 |
| **Z3** | 33,416,553 | 24,804,936 | 8,167,866 | 59,830,364 | 68,388 | 836 | 55.23 |
| **Z4** | 35,991,622 | 27,329,748 | 6,982,030 | 46,160,878 | 51,450 | 868 | 53.83 |
| **Uk1** | 31,718,066 | 25,270,230 | 15,346,535 | 97,468,932 | 93,004 | 1,075 | 85.81 |
| **Uk2** | 11,924,977 | 9,191,952 | 3,141,161 | 56,224,377 | 58,668 | 959 | 57.51 |

*Supplementary Table 3: Correlation between relative abundance of the top 20 bacterial orders and ARG load.*

| **Order** | **Mean relative abundance (%)** | **Spearman's correlation (relative abundance vs ARG load)** | |
|---|---|---|---|
| | | **Correlation coefficient** | ***p*-value** |
| **Aeromonadales** | 9.71 | 0.357 | 0.1916 |
| **Bacillales** | 2.36 | -0.304 | 0.2708 |
| **Burkholderiales** | 16.28 | 0.086 | 0.7630 |
| **Caulobacterales** | 1.22 | 0.243 | 0.3820 |
| **Corynebacteriales** | 3.29 | -0.439 | 0.1032 |
| **Deinococcales** | 1.09 | -0.104 | 0.7144 |
| **Enterobacterales** | 4.79 | -0.111 | 0.6953 |
| **Flavobacteriales** | 4.08 | 0.211 | 0.4499 |
| **Micrococcales** | 3.71 | -0.529 | 0.0454 * |
| **Micromonosporales** | 1.06 | -0.654 | 0.0100 * |
| **Nostocales** | 3.71 | -0.711 | 0.0041 * |
| **Propionibacteriales** | 1.45 | -0.664 | 0.0086 * |
| **Pseudomonadales** | 7.11 | 0.604 | 0.0195 * |
| **Rhizobiales** | 10.00 | -0.443 | 0.1002 |
| **Rhodobacterales** | 3.72 | -0.332 | 0.2264 |
| **Rhodocyclales** | 1.37 | -0.075 | 0.7926 |
| **Rhodospirillales** | 1.35 | -0.479 | 0.0735 |
| **Sphingomonadales** | 3.00 | -0.082 | 0.7728 |
| **Streptomycetales** | 3.22 | -0.661 | 0.0090 * |
| **Xanthomonadales** | 2.74 | -0.007 | 0.9847 |

** $p < 0.05$ = statistically significant.*

*Supplementary Figure 1. Species-level beta-diversity: a) Bray-Curtis dissimilarity heatmap; b) Principal coordinates analysis (PCoA) performed using Bray-Curtis distances.*

**a**

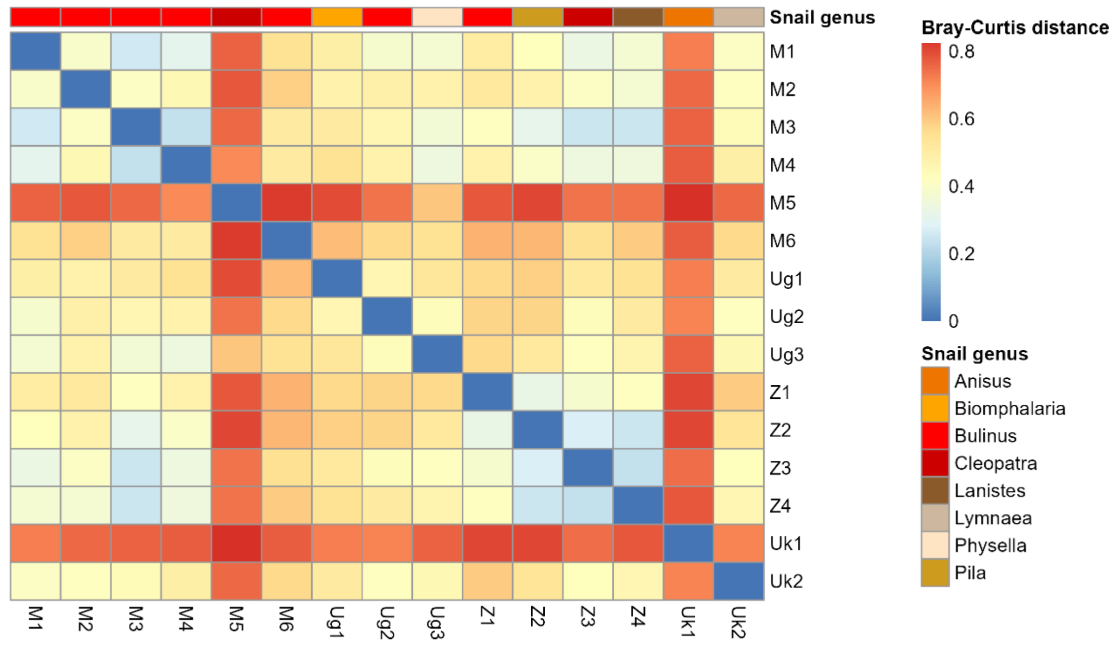


**b**

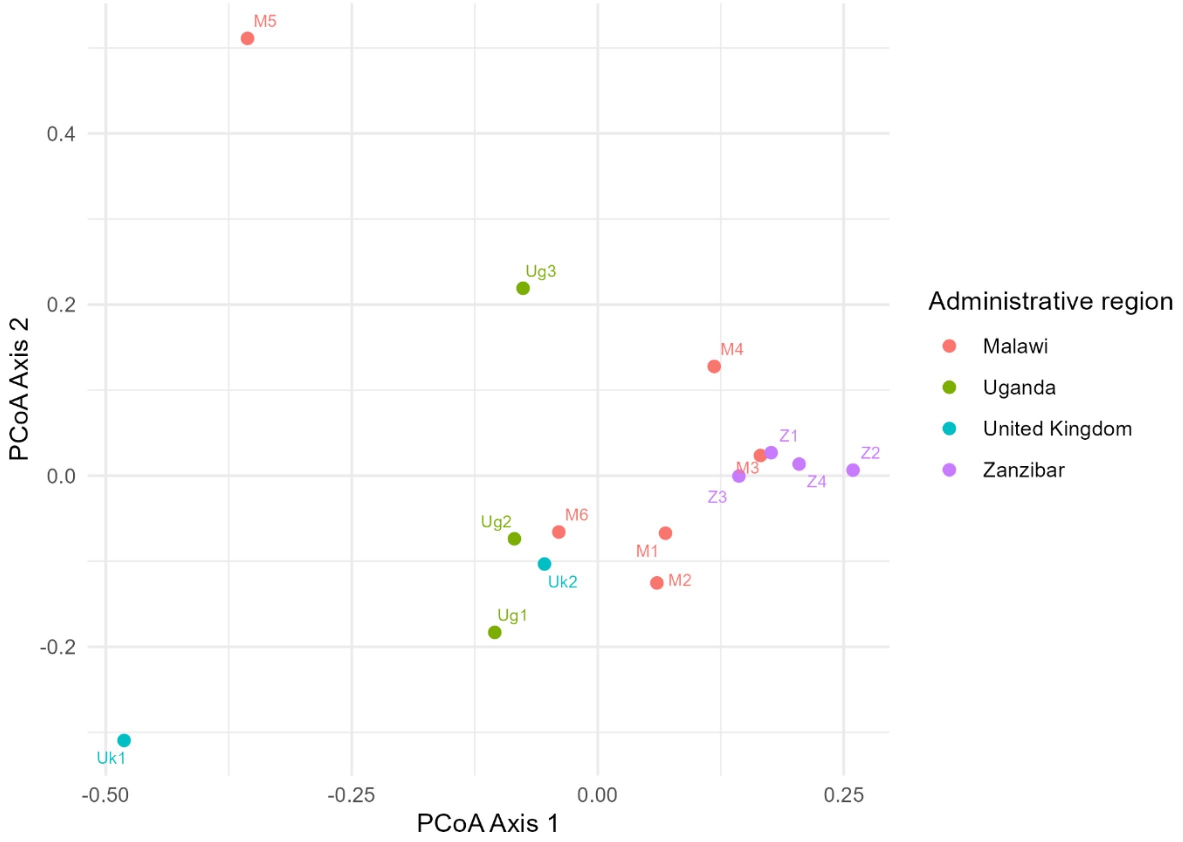

*Supplementary Figure 2: Scatter plots showing associations of sequencing depth with: a) ARG detection; b) MAG recovery.*

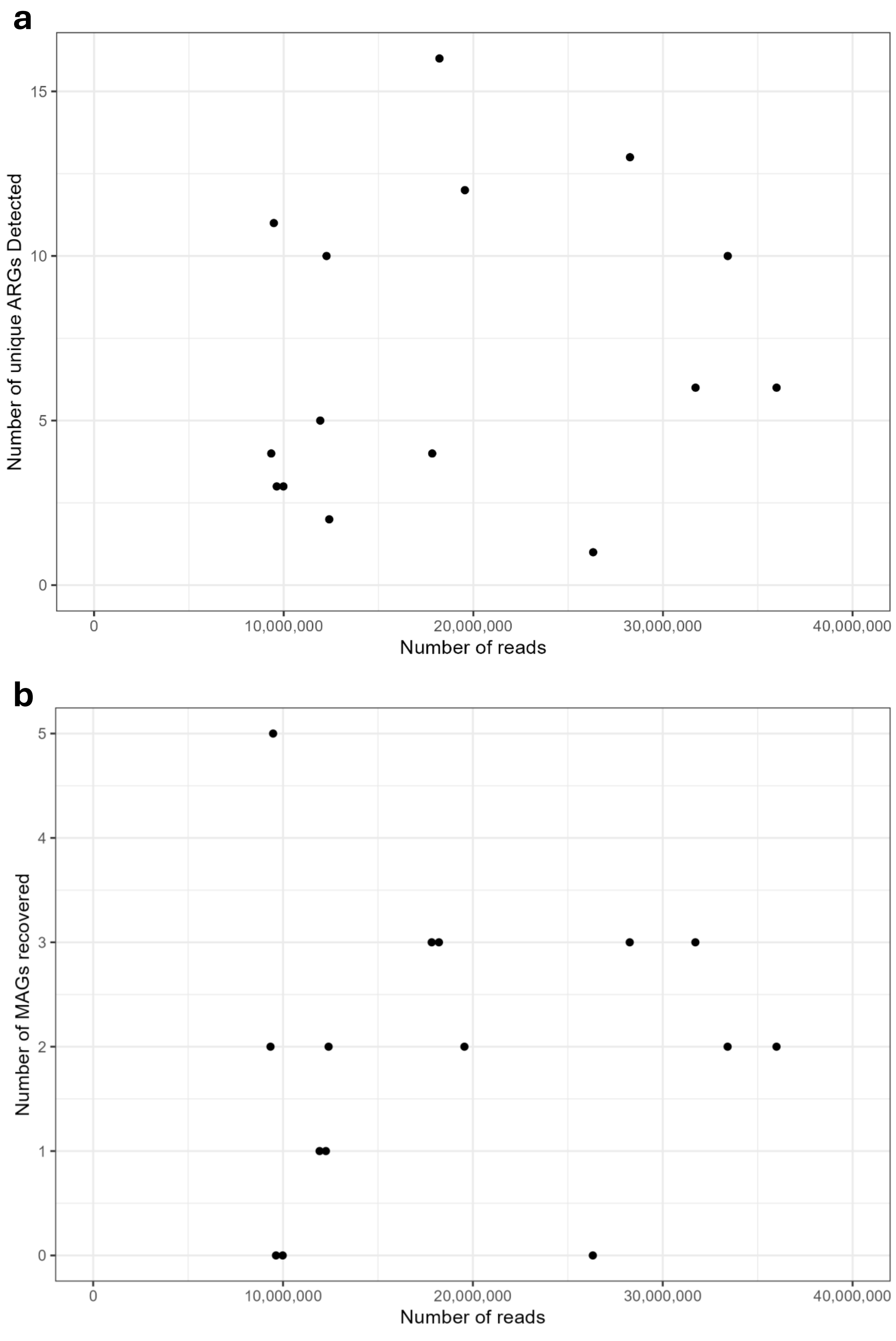

*Supplementary Figure 3: Rarefaction curves showing the number of unique ARGs detected following random sub-setting of reads from samples with > 20 million reads following trimming and filtering.*

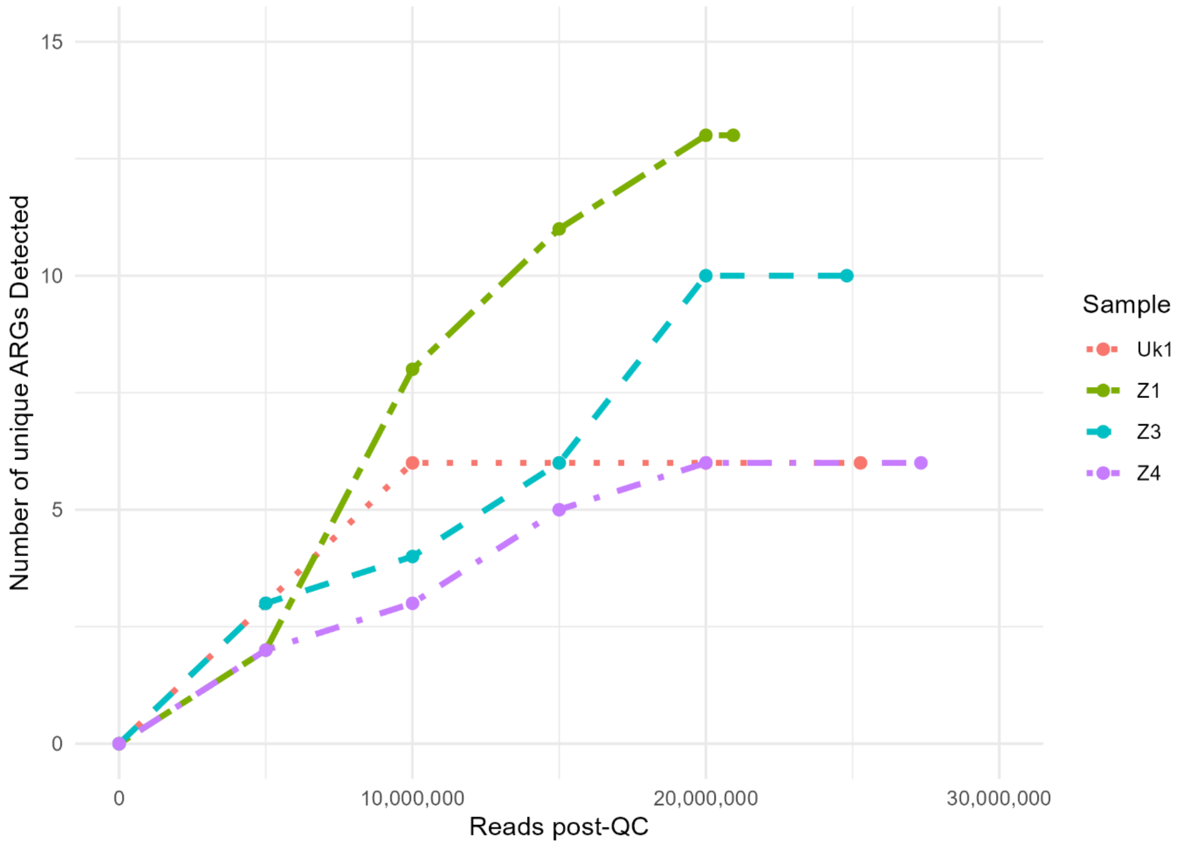

*Supplementary Figure 4: Relative abundance of ARGs aggregated by class (RPKM).*

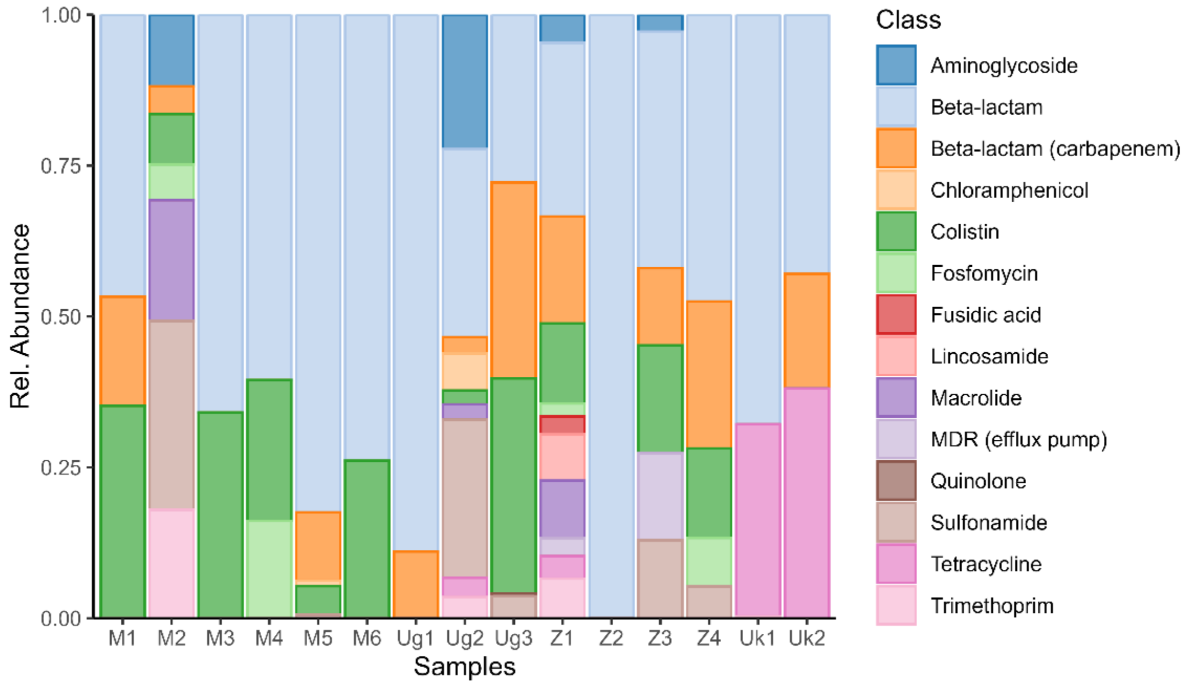

*Supplementary Figure 5: Read- versus assembly-based ARG detection.*

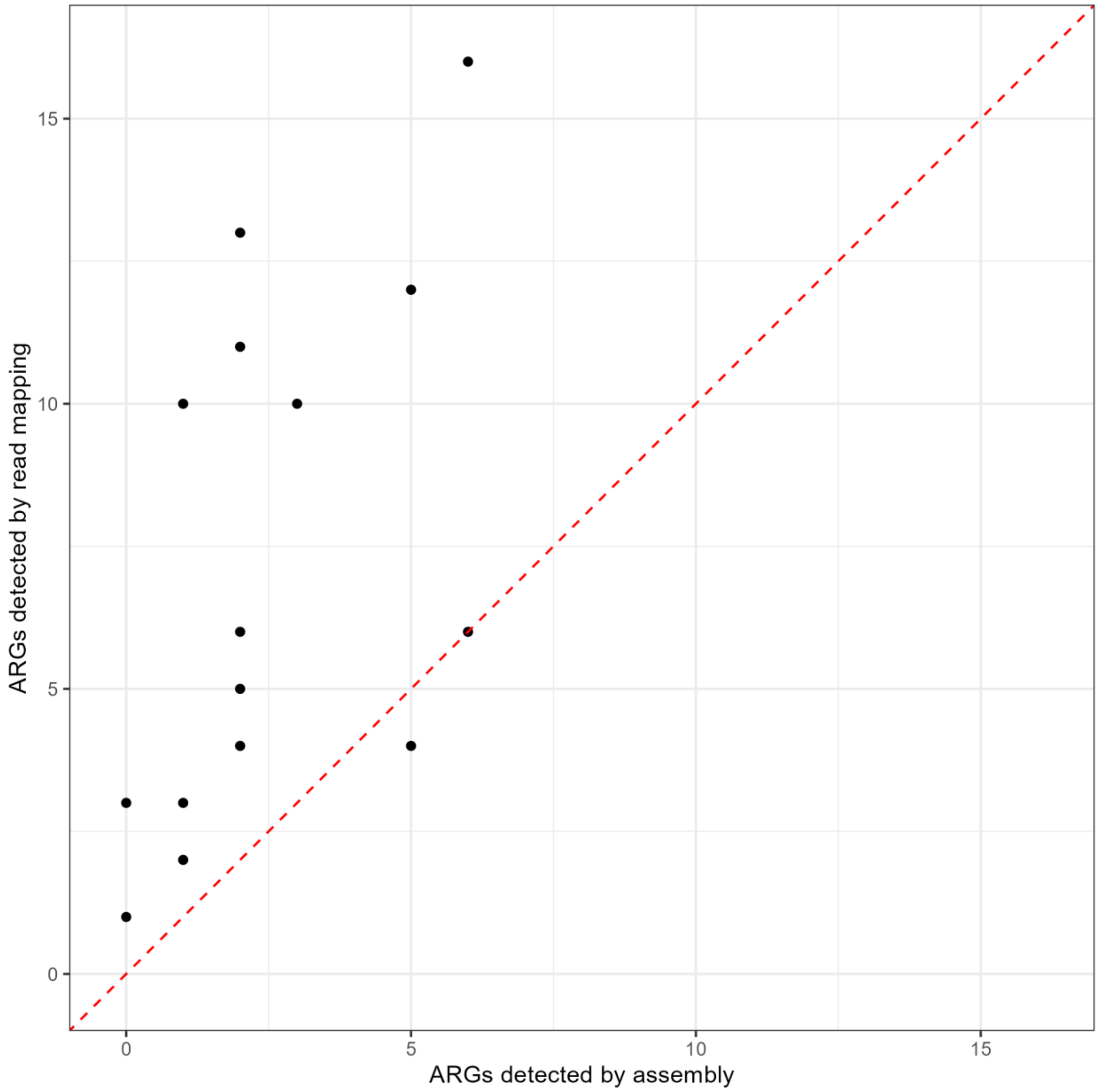

*Supplementary Figure 6: Annotation of M5 NODE_173 containing bla$_{OXA-181}$ and ΔlysR, compared to Tn2013 from K. pneumoniae strain KP3 plasmid pKP3-A (Accession no. JN205800) containing bla$_{OXA-181}$ flanked by ΔlysR-ΔereA and ISEcp1.*

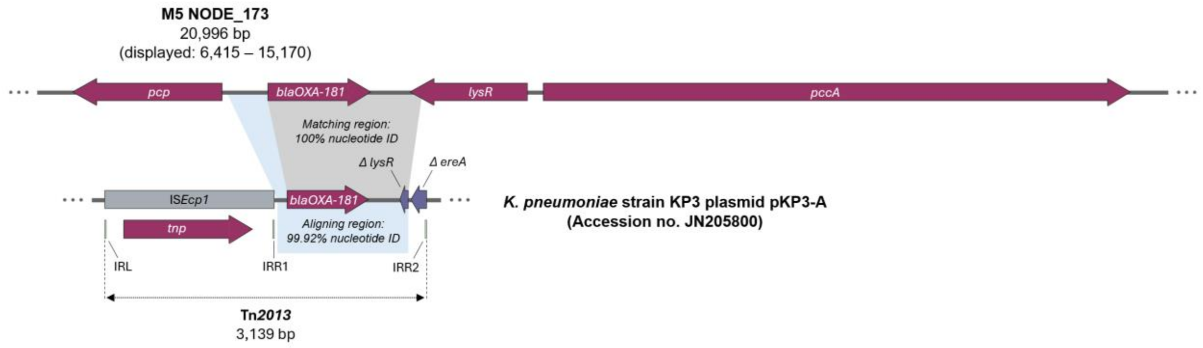
